# A Novel Phenotyping Approach for Reconciling Precision and Variance in Disease Severity Estimates from High-resolution Imaging

**DOI:** 10.64898/2026.02.20.707028

**Authors:** Radek Zenkl, Bruce A. McDonald, Jonas Anderegg

## Abstract

Accurate quantification of plant disease is essential for resistance breeding, variety testing, and precision agriculture, yet visual ratings are limited by subjectivity, low precision, and restricted throughput. Image-based phenotyping can address these limitations, but field applications face substantial challenges due to spatial heterogeneity, symptom-level diagnostic requirements, and the need for very high-resolution imagery with limited spatial coverage. This introduces a fundamental trade-off: high-resolution images provide precise local measurements of disease, but spot-level estimates can be highly variable within experimental units.

We analyzed a large image data set of wheat foliar diseases to characterize the distribution, spatial dependence, and aggregation behavior of spot-level severity estimates in plots. We combined high-resolution macro-scale imaging with focus bracketing to increase the sampled leaf area. Our results highlight focus bracketing as a promising approach for simultaneous diagnosis and quantification of disease in field plots. Autocorrelation in severity estimates both within focal image stacks and across plot positions was comparable, with 10 focal stack images or 10 positions per plot contributing approximately 2.5 independent observations each. Modeling plot-level severity as a latent Beta-distributed variable enabled robust estimation of mean severity and associated uncertainty. This supports both hypothesis testing and efficient sampling across the full range of disease severity associated with genotypic diversity and seasonality of developing epidemics.

The proposed imaging approach is non-invasive and, in principle, transferrable to autonomous ground-based phenotyping platforms, offering the potential to shift the dominant source of uncertainty in estimating disease severity from measurement-related limitations toward biologically and environmentally driven variability in disease expression.

## 2 Introduction

Accurate and precise measurement of plant disease is essential for diverse applications, ranging from fungicide development to resistance breeding, and from elucidating molecular host–pathogen interactions to understanding epidemiological processes at the landscape scale [6]. In most contexts, visual rating of disease symptoms by trained experts remains the gold standard for detection and diagnosis as well as for quantitative evaluation of severity [9]. However, visual ratings are constrained by limited precision, accuracy, and throughput, as well as by the level of detail that can be achieved in terms of symptom characterization [2, 7, 23, 35]. These limitations have prompted increasing interest in sensor-based approaches that may yield perfectly reproducible measurements of many aspects of disease, thus eliminating the problem of intra-rater variability. Given that the same method can be applied consistently across experiments, mimicking a situation in which one rater accomplishes all phenotyping work, inter-rater variability (i.e., rater bias) may also be eliminated [7].

Image-based approaches have shown strong potential to improve accuracy and precision of severity assessments, particularly for organ-level assessments under controlled conditions. In such assays, several factors simplify the task of obtaining reliable image-based estimates: i) inoculations are often made with a single or a few pathogen strains on seedling leaves, reducing symptom phenotypic variability; ii) imaging setups and conditions such as lighting can be tightly controlled, minimizing scene complexity and variability; iii) other stress symptoms are usually absent, eliminating the need for symptom-level diagnosis; and iv) the entire experimental unit can often be imaged at once and in a standardized manner, eliminating the need for aggregation of partial measurements to the scale of the experimental unit.

Due to these favorable circumstances, relatively straight-forward analytical image analysis approaches may be sufficient for direct and accurate trait extraction from single images (e.g., [5, 12, 13, 22, 29, 34]). Controlled environment studies have been invaluable for advancing our understanding of plant–pathogen interactions, yet field evaluations are ultimately required for practical application. In the field, however, the simplifying conditions of controlled environments no longer apply: inoculum sources and host morphologies are diverse, environmental conditions vary, and other stresses are common. As a result, accurate disease quantification often requires diagnosis at the symptom level in complex and variable scenarios. Some representative examples include: 1) the need to distinguish necrosis occurring at the leaf tip in wheat cultivars that possess the quantitative resistance gene *Lr34* from necrosis caused by necrotrophic pathogens; 2) the need to separate necrotic lesions caused by Septoria tritici blotch (caused by *Zymoseptoria tritici*) from those caused by Septoria nodorum blotch (caused by *Parastagonospora nodorum*) on wheat leaves, which often co-occur. If these challenges are not adequately addressed, the previously mentioned rater bias would simply be replaced by biases related to automated image interpretation, introducing new sources of error that could have been effectively handled by experienced human raters.

The requirement for symptom-level diagnosis, in turn, demands very high-quality imagery with fine spatial resolution, since diagnostic features that enable such distinctions are often extremely small. For example, the black pycnidia (i.e., fruiting bodies containing asexually produced spores) produced by *Z. tritici* within necrotic lesions used for symptom diagnosis are well below 1 mm in diameter, and imagery with a resolution of approximately 0.02 mm / pixel is necessary to be able annotate and reliably detect them [33, 40]. Approaches using lower resolution imagery have been explored for disease detection and quantification (e.g., [3, 1, 10]), but these are typically restricted to anomaly detection or overall plant health monitoring and currently offer very limited potential for accurate diagnosis [3, 24, 25].

The high requirements in terms of image quality mean that a single measurement rarely represents a significant portion of an experimental unit in field experiments, which may consist of an infection row or a microplot in breeding trials, or larger plots in variety testing trials. In fact, not even a single plant or plant organ may be fully represented in one image, due to the small image foot print and shallow depth-of-field inherent to close-up imaging, combined with uncontrolled positioning and orientation of plant organs with respect to the focal plane (see e.g., [39]). As a result, using high-resolution images for disease assessments creates a tradeoff between precision and variance, where each image yields an accurate and precise representation of a particular spot, but the spot-level measurements introduce a high variance into the estimates of the overall disease level in a plot. This problem is particularly pronounced for diseases that are caused by soil- or residue-borne and smear- or splash-dispersed pathogens such as *Z. tritici* or *P. nodorum* on wheat, or *Phytophthora* spp. on various crops, which tend to create spatially heterogeneous disease severity due to local spread (e.g., [17]). Additional heterogeneity may arise from plot border effects, microclimatic variation, differences in canopy structure, and uneven initial inoculum exposure.

These challenges are not unique to sensor-based phenotyping but also affect traditional, visual assessments. In organ-level evaluations, a rater can typically view the entire experimental unit at once, which is impossible in field plots. Instead, plot-level estimates often rely on aggregating organ-level observations. For example, a visual rating of disease incidence (the fraction of infected plants or organs) can be combined with severity ratings on infected leaves to provide an accurate estimate of overall plot-level disease [1, 4]. Such approaches can even take the localization of symptoms within the canopy explicitly into account when quantifying overall damage (e.g., [11, 20]). However, such detailed assessments are far more time-consuming than a single, integrated ‘snapshot’ rating of overall disease severity at the plot level, because sample sizes must be large to achieve sufficient precision.

When multiple diseases must be monitored across hundreds or thousands of experimental units, often replicated across different environments, a rater typically has only seconds to evaluate each unit. Under such time constraints, the rater has to aggregate multiple observations implicitly, and this complex cognitive process may represent a major source for both intra- and interrater variability. Notably, whereas errors in visual organ-level estimates of disease severity have been well studied (reviewed by Bock et al. [7]) these plot-level aspects of variability have received very little attention [27, 28].

Novel image-based approaches to disease quantification offer the opportunity to more explicitly aggregate spot-specific measurements to the level of the experimental unit, similar to what can be achieved using time-consuming manual scoring protocols that rely on explicit organ-level assessments [32]. This is particularly the case for measurement procedures that do not require physical interaction with plants and are therefore amenable to automated data acquisition. Despite this promise, the use of multiple measurements per experimental unit to improve precision remains rare.

Here, we leverage a large, high-resolution image data set comprising 16,650 images of 15 wheat plots taken on two consecutive days to investigate the statistical distribution and within-plot spatial dependencies of disease-related trait values. We combine a previously developed imaging protocol and image processing workflow [39] with focus bracketing to compensate for the shallow depth-of-field resulting from macro photography in order to increase the analyzed leaf area. The resulting unique data set allows us to address three important questions: (i) what is a typical distribution of disease severity metrics estimated from high-resolution images that each represent a specific spot in an experimental plot? (ii) how many spot-level measurements are required to reach a desired level of precision? and (iii) what is the best way to aggregate spot-level measurements into reliable plot-level estimates, given that severity scores are inherently bound between 0 and 1 and individual estimates from repeated measurements are auto-correlated? Our analysis demonstrates the value of focus bracketing as a way of increasing the sampled leaf area at a negligible marginal cost. It also provides novel insights into the distribution and variability of estimated spot-level disease severity estimates, and leads to a practical approach for aggregating spot-level measurements at the level of plots as common experimental units.

## 3 Materials and Methods

### 3.1 Field Experiment and Data Acquisition

A field experiment was conducted at the ETH Research Station for Plant Sciences in Lindau-Eschikon, Switzerland (47.449°N, 8.682°E), during the 2022–2023 wheat growing season. The experiment included 16 wheat cultivars selected based on their contrasting morphology and resistance to STB, with the aim of including a diverse set of traits in terms of canopy structure, plant morphology, and STB symptom expression. Each cultivar was grown in nine microplots sized 1 m x 1.7 m, of which three were assigned each to one of the following treatments: i) an early fungicide application with a later artificial inoculation with a blastospore solution containing 10 *Z. tritici* isolates; ii) an early fungicide application without subsequent inoculation; and iii) no fungicide or inoculation treatment. These treatments generated a broad range of disease symptoms and severities from both natural and artificial infections (for more details on experimental design, see [3]).

High-resolution RGB images were collected following a protocol described previously [39], with some modifications. Briefly, imaging was performed using a hand-held full-frame mirrorless digital camera (EOS R5, Canon Inc. Tokyo, Japan; 45 megapixel, 36 x 24 mm sensor) combined with a macro lens (RF 35 mm f/1.8 IS Macro STM; Canon Inc., Tokyo, Japan). A custom-made acrylic glass frame was attached to the camera lens and used as a gauge to ensure a consistent distance of 18 cm between the camera and the outermost parts of the canopy. This imaging setup resulted in a sampling distance of 0.022 mm / px. The resulting field of view was approximately 11 × 17cm.

At each imaging position, a stack of images was acquired using focus bracketing mode. This technique captures a sequence of images while systematically shifting the focal plane, producing a focal stack of many images where different regions are in focus (Figure 1). The digital camera used in this study automates this process, capturing the entire stack of images with a single button press. The focus distance is incrementally adjusted in predefined steps that were set to ensure some overlap between the in-focus regions of consecutive frames. A focal stack of ten images was captured at each imaging position, taking approximately 1 second. The depth-of-field in the first image of the stack was approximately 5 mm, as experimentally determined. The depth-of-field progressively increases along the stack as does the asymmetry of the depth-of-field, such that the proportion on the far side relative to the near side increases. The final frame was set to maintain sufficient resolution for imaging of small features, such as pycnidia. A resolution of approximately 0.026 mm / px at a distance of 20.5 cm, corresponding to the distance of the final focal plane, was achieved. The distance between the initial and the final focal planes amounted to approximately 25 mm, while the total distance from the nearest to the farthest point in acceptable focus exceeded 30 mm. Given the side-view imaging geometry and the high density of wheat stands, the total depth-of-field achieved was sufficient to capture most of the relevant exposed tissues in focus in at least one frame. The camera retains the final focus setting after capture of a focal stack, which requires refocusing to a known reference point representing the nearest focal plane, which was achieved using the aforementioned distance gauge.

**Figure 1.**
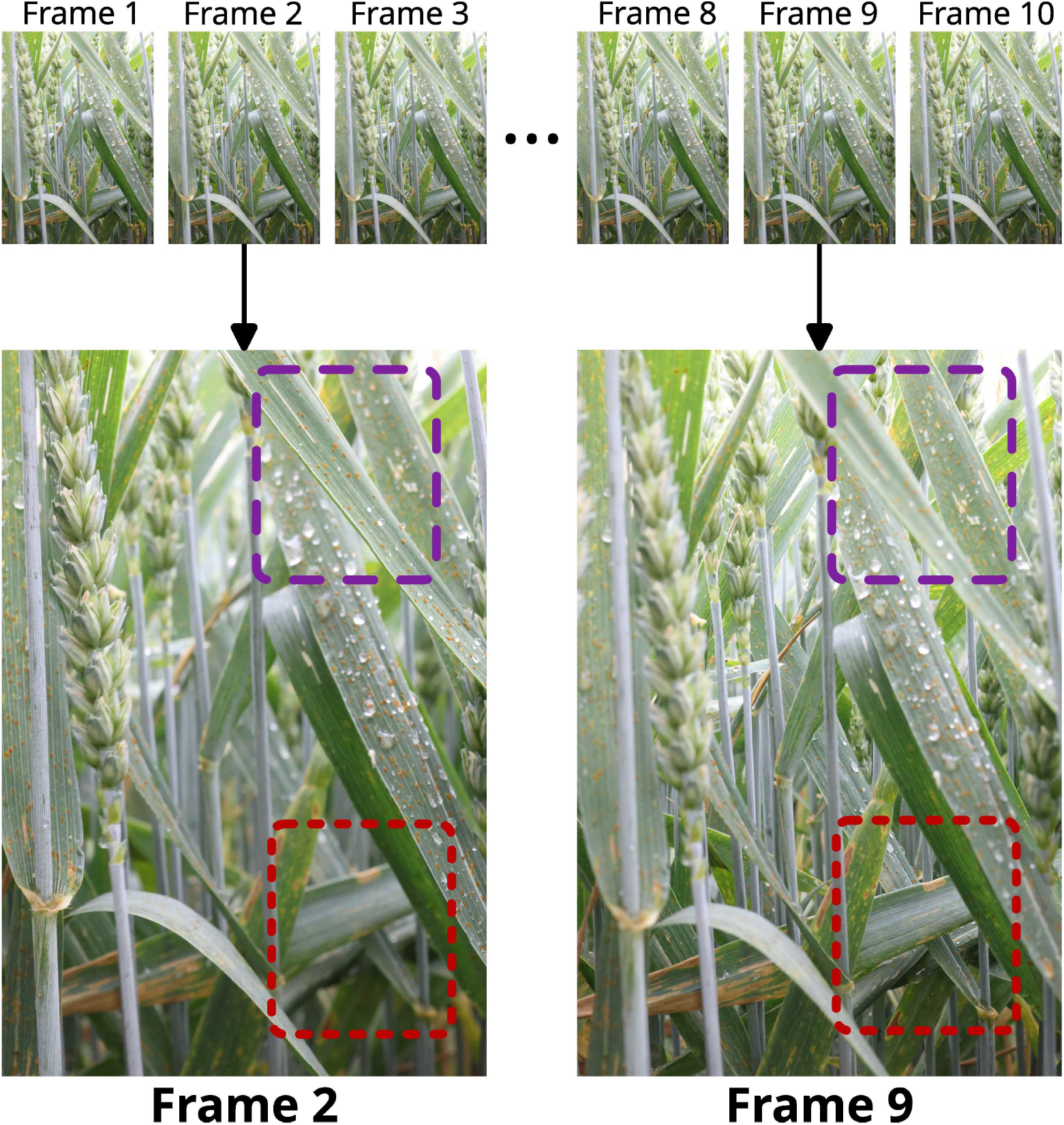
Visualization of a focal stack consisting of ten frames, each captured with a slightly different focal plane. This variation is illustrated in the areas highlighted by the purple and red rectangles. In the purple rectangle, the front leaf is in focus in frame 2 but becomes blurry in frame 9. In contrast, the leaf in the red rectangle appears blurry in frame 2 but comes into focus in frame 9. These differences demonstrate how shifting the focal plane across the frames affects which parts of the scene appear sharp or blurred.

Here, a subset of 16,650 images taken in 15 untreated plots on June 6 and 7, 2023 was used for in-depth analyses of the distribution of extracted trait values and auto-correlation patterns resulting from multiple plot measurements. The plots were extensively imaged from approximately 32 different locations along all sides of the plot, moving systematically to the next location around the plot. This resulted in a pitch of approximately 14 cm. At each position, the imaging setup was aligned with the top three leaf layers, starting with the flag leaf, producing 32 focal stacks for each of the top three leaf layers.

### 3.2 Trait extraction from images

Disease severity measures were extracted from images using deep-learning-based image segmentation and keypoint detection as per our previous work [40, 39]. Briefly, in-focus leaf area was extracted using organ segmentation and a combination of texture analysis and depth estimation to identify leaf areas with sufficient quality for unbiased disease evaluation. Rust pustules and necrotic lesions were detected and necrotic lesions segmented using specifically trained Yolo v11 [16] and Segformer backbone [37] with FPN head [21], respectively. So far, a distinction between yellow rust and brown rust pustules is not made, but brown rust was overwhelmingly dominant at the time of image acquisition. Based on the resulting masks, percent leaf area covered by lesions (PLACL) and rust pustule density were extracted for each image for further analyses, using the total in-focus leaf area as the corresponding reference.

### 3.3 Modeling Distributions of Concurrent Severity Estimates at the Plot Level

Accurately modeling individual PLACL measurements as realizations of a latent random variable enables deeper and more rigorous statistical analyses. A well-specified distribution allows for the estimation of confidence intervals and quantification of uncertainty.

PLACL represents the proportion of diseased leaf area, making it naturally bounded within the interval [0, 1]. Such fractional data cannot be appropriately modeled using unbounded distributions, as these may predict biologically meaningless values outside this range. To address this, we employed the Beta distribution, a continuous probability distribution defined by two shape parameters,*X* ~ Beta(*α, β*). The Beta distribution can represent a wide range of shapes, including uniform, unimodal, and skewed forms (see Figure 2).

**Figure 2.**
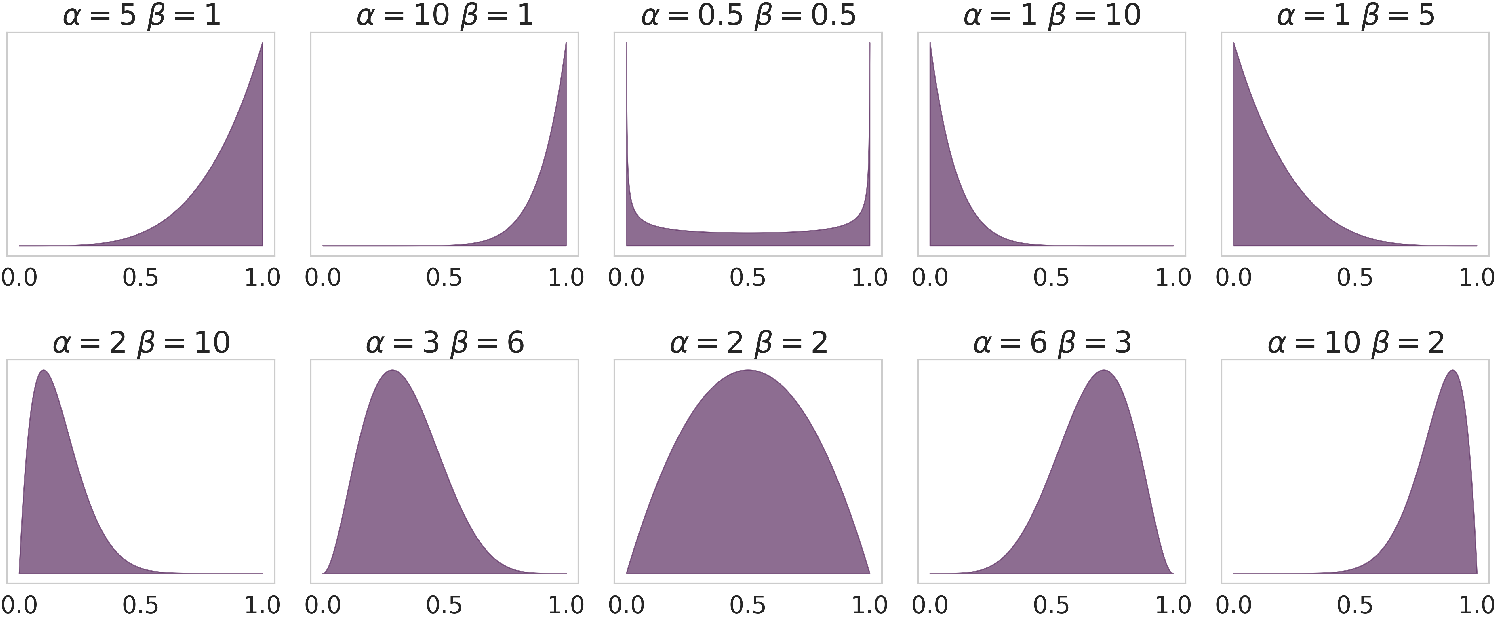
Resulting shapes of Beta(*α, β*) distribution based on different values of *α* and *β*.

**Figure 3.**
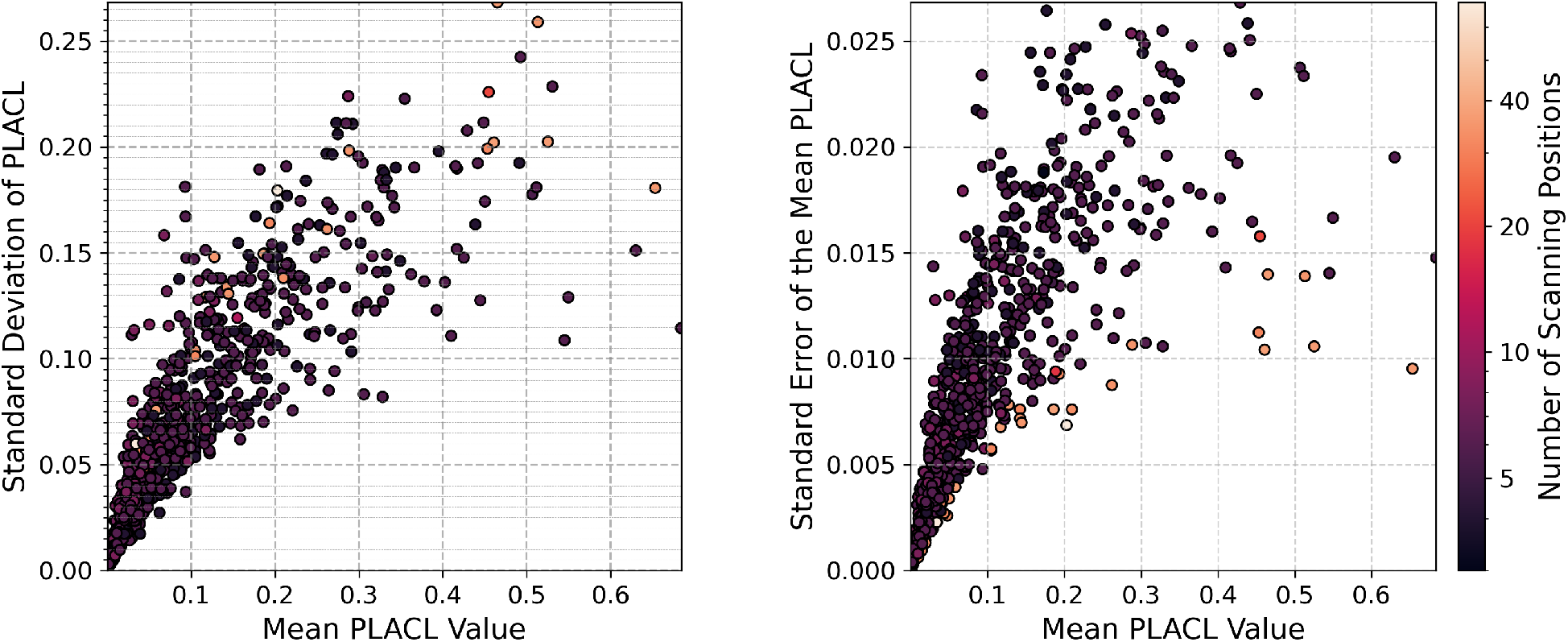
Standard Deviation (STD) of PLACL and Standard Error of the Mean (SEM) of PLACL plotted as a function of mean PLACL. The number of scanning positions around the plot is denoted in form of the color of the plotted data points.

This flexibility makes it particularly well-suited for modeling PLACL. In field experiments, disease severity often shows substantial heterogeneity across plots, ranging from nearly healthy to highly diseased canopies, resulting in distributions that may be skewed. The Beta distribution can capture these patterns while providing interpretable parameters such as its mean which represents the expected severity, and its variance reflecting the uncertainty or heterogeneity among measurements.

### 3.4 Autocorrelation in focal stacks and across multiple positions

All images in a focal stack capture the same area of a plot, depicting the same plants and plant organs. Because consecutive images in the stack share partially overlapping focal planes, there is partial overlap in the represented plant material, necessarily leading to correlated measurements within a stack. Similarly, measurements taken at different positions within a plot may also be correlated, as consecutive positions capture nearby plants or plant organs, possibly representing the same disease foci, even though the exact same plant material is not present across positions.

To assess the degree of non-independence of trait values derived from images within a focal stack, we estimated their autocorrelation. This was first done for each focal stack separately by estimating the sample autocorrelation function (ACF) of trait values up to lag *k* =9 using the base R function *acf()*. This approach estimates the correlation between the trait values at position *t* and *t + k*, standardized by the variance of the series, across all pairs available within the stack. Due to the small sample size, particularly at larger lags, the resulting estimates are inherently noisy and subject to uncertainty. To obtain a more reliable overall picture of autocorrelation patterns, we summarized these estimates by calculating the mean and standard deviation of the ACF at each lag. We explored potential effects of genotype, leaf layer, and disease severity by summarizing trends at respective grouping levels. This allowed us to visualize the typical decay of the correlation with increasing distance between images in a stack while also capturing the variability between stacks.

A similar procedure was implemented to assess non-independence of trait values derived from focal stacks captured at different positions within plots. For this, trait values were first averaged across all images in a focal stack and the sample ACF up to lag 24 was estimated as described above. As for focal stacks, potential effects of leaf layer and genotype characteristics were explored by summarizing trends at the respective grouping levels.

To formally model this correlation structure using the entire data set, we fitted a linear mixed-effects model with a random intercept for each focal stack and an auto-regressive correlation structure of order 1 (AR(1)) along the images in the stack, using the R function *lme()* of the R-package ‘nlme’. This model-based approach provides a single estimate of the within-stack serial correlation while controlling for potential plot- and leaf-layer effects. The AR(1) auto-regressive correlation structure was indicated by the apparent exponential decay of the autocorrelation estimated for each stack independently. The fitted model was

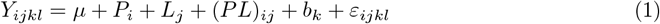

where *Y*_*ijkl*_ is the trait value for image *l* in stack *k*, leaf layer *j*, and plot *i*; *µ* is the overall intercept; *P*_*i*_ is the fixed effect of plot (genotype) *i*; *L*_*j*_ is the fixed effect of leaf layer *j*; (*PL*)_*ij*_ is the interaction between plot and leaf layer; *b*_*k*_ is the random effect of stack *k*; and *ε*_*ijkl*_ is the residual error, assumed to follow an AR(1) correlation along the sequence of images within each stack. Each plot was measured at multiple positions attributed a numeric index (1… 32). Measurement position within plots was thus specified as nested in plot in the random term. PLACL data were logit-transformed, pustule density data were log-transformed prior to modeling. Model assumptions were evaluated by inspecting residual plots, including fitted values versus residuals, Q–Q plots, and histograms of residuals to assess normality and homoscedasticity (Supplementary Figure S2).

To assess the degree of non-independence of trait values derived from focal stacks captured at different positions within a plot, we fitted linear mixed models with an AR(1) structure to the sequence of positions. We first obtained fitted values from the stack-level mixed model (accounting for within-stack correlation; eq. 1), and then averaged these fitted values at the stack level. These position-level summaries were analyzed with plot, leaf layer, and their interaction as fixed effects. Random intercepts were included for each plot-by-layer combination to account for baseline differences in trait levels. The AR(1) structure was specified within each plot-by-layer combination. Thus, the model was

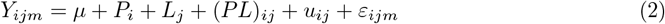

where *Y*_*ijm*_ is the stack-level fitted trait value at position *m*, leaf layer *j*, and plot *i*; *µ* is the overall intercept; *P*_*i*_ is the fixed effect of plot (genotype) *i*; *L*_*j*_ is the fixed effect of leaf layer *j*; (*PL*)_*ij*_ is the interaction effect; *u*_*ij*_ ~ 𝒩 (0, *σ*^2^) is a random intercept for each plot-by-layer combination; and *ε*_*ijm*_ is the residual error, assumed to follow an AR(1) correlation along position within each plot-by-layer combination. Models including fixed-factor interactions were preferred over models without interactions based on a comparison of nested models using the Akaike Information Criterion (AIC). More complex serial correlation structures (e.g., ARMA models) yielded marginally lower AIC scores, but the simpler AR(1) structure was finally preferred because it provides a clear representation of serial autocorrelation, corresponding to an exponential decay of correlation with increasing image distance. The AR(1) model also requires fewer parameters to be estimated, reducing the risk of overfitting given the limited number of observations per image sequence.

The estimated autocorrelation coefficient (*ϕ*_1_) quantifies autocorrelation on the transformed scale. However, from a breeder’s perspective, the original scale (i.e., absolute PLACL and rust pustule density) is more relevant. To obtain an estimate of autocorrelation comparable to the sample-based estimate, we back-transformed simulated trait values from the fitted model to the original scale and then computed per-stack autocorrelation, averaging across stacks for a robust, model-based estimate.

Estimated sample-based and model-based acf were then used to estimate the effective sample sizes resulting from measurements across multiple positions-in-plot and positions-in-stack, based on the following equation;

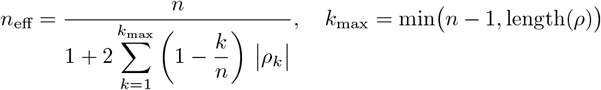

where *n* is the total number of observations and *ρ*_*k*_ is the estimated autocorrelation at lag *k*. To enable a direct comparison of estimated effective sample sizes across groups with varying number of measurements, the effective sample size was divided by the total number of samples to obtain the effective sample fraction (ESF):

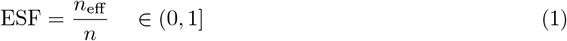

This approach follows the general principle of variance inflation due to autocorrelation and has been applied in ecological research [31].

R-code to reproduce the full analysis is available at https://github.com/and-jonas/plot-spot-analysis.

## 4 Results

### 4.1 Modeling Plot-level PLACL as a Beta-Distributed Trait

To empirically test the suitability of the Beta distribution to represent the observed distribution of estimated trait values, it was fitted to data from the most extensive imaging campaign, in which 15 different cultivars were imaged (see Section 3.1). The fitted Beta distributions are shown alongside the corresponding trait value histograms in Figure 4. Overall, the Beta distribution provided a visually convincing fit across all plots.

**Figure 4.**
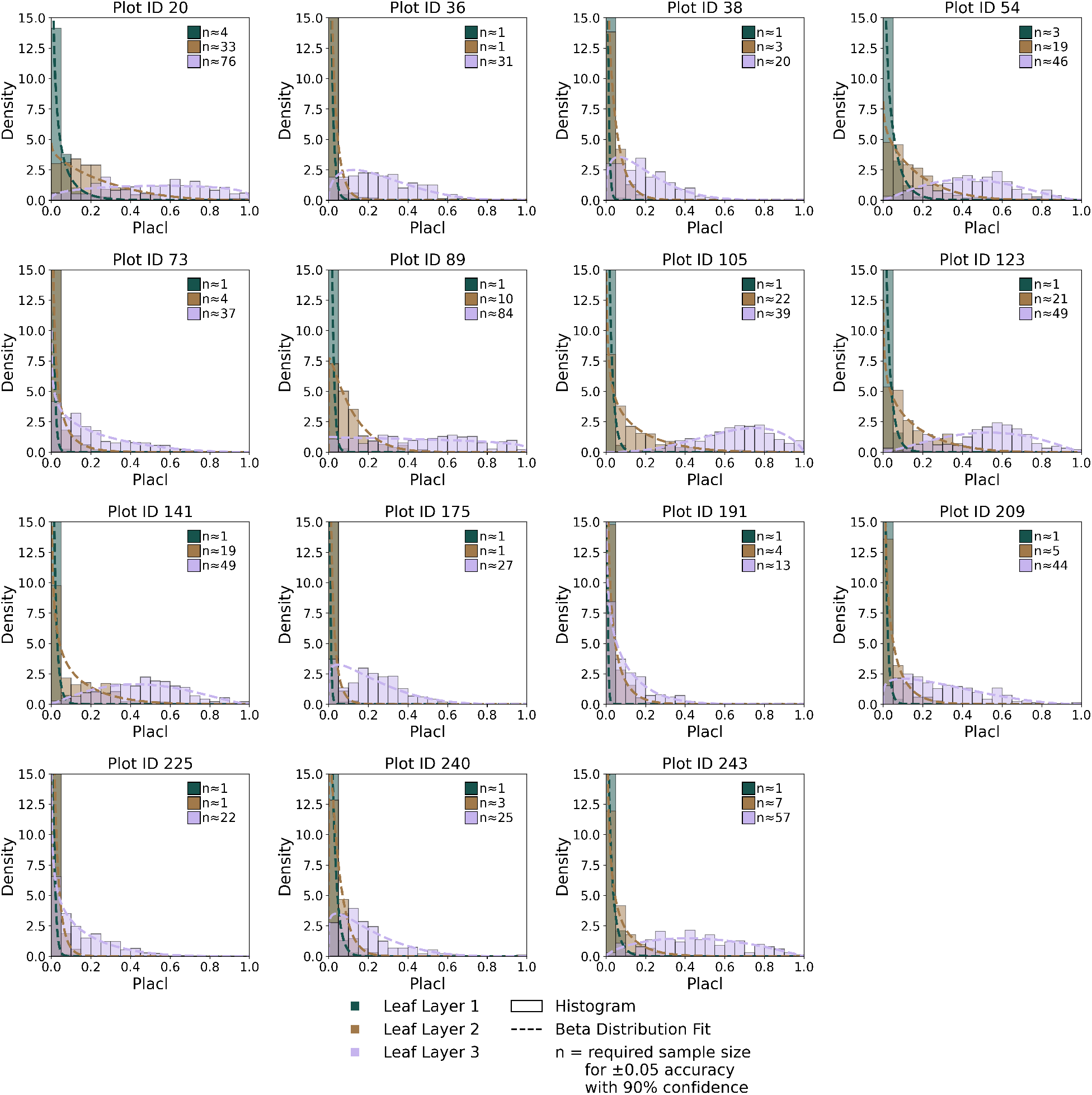
Fitted Beta(*α, β*) distributions for various cultivars and leaf layers. Fitted Beta(*α, β*) is denoted with a dashed line. The raw measurement data is visualized using a histogram. Layer 1 corresponds to the flag leaf layer. The x-axis represents PLACL values in the range [0,1], and the y-axis represents the density of the histogram and the corresponding Beta distribution.

There was a general trend for the average severity density to decrease when moving up the canopy (Figure 4), which is in line with expectations based on the soil-borne and splash-dispersed nature of the pathogen. STB severity estimates for Layers 1 and 2 were heavily left-skewed (i.e., zero-inflated-like). Strongly contrasting severity distributions across plots were observed particularly for layer 3, with fitted Beta distributions ranging from strongly left-skewed (zero-inflated–like), to approximately symmetric, and to right-skewed (but not one-saturated) forms. This reflected substantial variation in the distribution of trait values between cultivars (plots). Several plots exhibited a mean severity close to 0.5, but differed markedly in the shape of their distributions (e.g., PlotID 89 vs. PlotID 123; Figure 4). Some exhibited relatively symmetric distributions (*α* ≈ *β* ≈ 1), while others exhibited peaked or skewed profiles. These differences in distribution reflect spatial variation in disease prevalence and severity within the canopy, potentially offering insights into plot-level disease dynamics beyond mean severity.

To estimate the parameters underlying the Beta distribution with a given precision and confidence, the degree of uniformity of the data needs to be taken into account. For example, depending on the shape of the distribution, between 1 and 84 samples are required to estimate the mean within *±*0.05 with a probability of 90% (Figure 4).

### 4.2 Increasing the Analyzed Leaf Area with Focus Bracketing

On average, 19.7, cm^2^ of leaf area was evaluated in each image, which exceeds the average leaf size used for evaluations in [19] (see Section 2). On average, approximately 23% of an image was predicted to be in focus, with this fraction increasing with increasing focal distance (Figure 5). Thus, as expected based on the imaging setup and procedure, there was a sizeable overlap between focused regions in subsequent images, leading to analyzing the same physical area multiple times. Assuming a static scene during image acquisition, an ideal focal stack could capture the scene so that everything is in focus at some frame in the stack. However, a more realistic estimate is that about 90% of the scene is relevant as the canopy is not completely dense in most cases, meaning that some irrelevant background, such as neighboring plots, sky or soil, can be visible. Following this estimate and considering the average fraction in focus per image, a theoretical maximum for increasing the sample size by utilizing focal stacks is given as:

**Figure 5.**
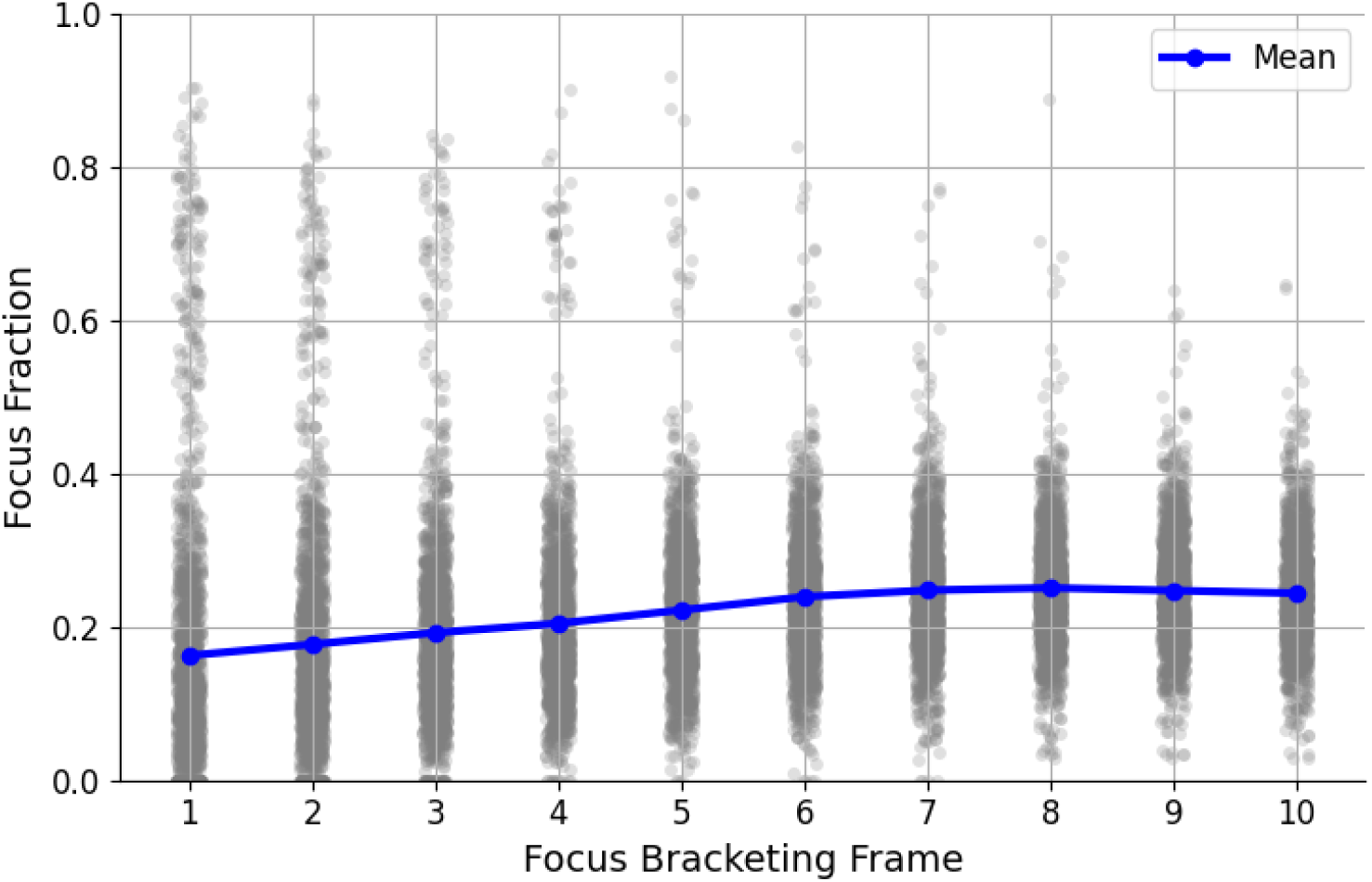
Ratio of the image predicted to be in focus plotted as a function of the position within a focal stack.

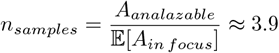

indicating that 10 images of a focal stack would provide the equivalent of 3.9 independent measurements. However, this estimate does not consider autocorrelation in the data related to spatial proximity, rather than related to overlap of leaf areas analyzed,which further reduces the effective sample size.

### 4.3 Autocorrelation patterns in image stacks and across imaging positions

Initial images of focal stacks generally yielded lower estimates of PLACL and rust pustule density than terminal images (Figure 6A). This pattern was quite consistent across leaf layers for rust pustule density, whereas for PLACL it was more pronounced at lower leaf layers corresponding to a higher severity. Thus, autocorrelation across focal stacks arose not only from overlapping plant material in consecutive focal planes and the same plants or plant organs being represented, but also from systematic trends in PLACL and rust pustule density along the stack.

**Figure 6.**
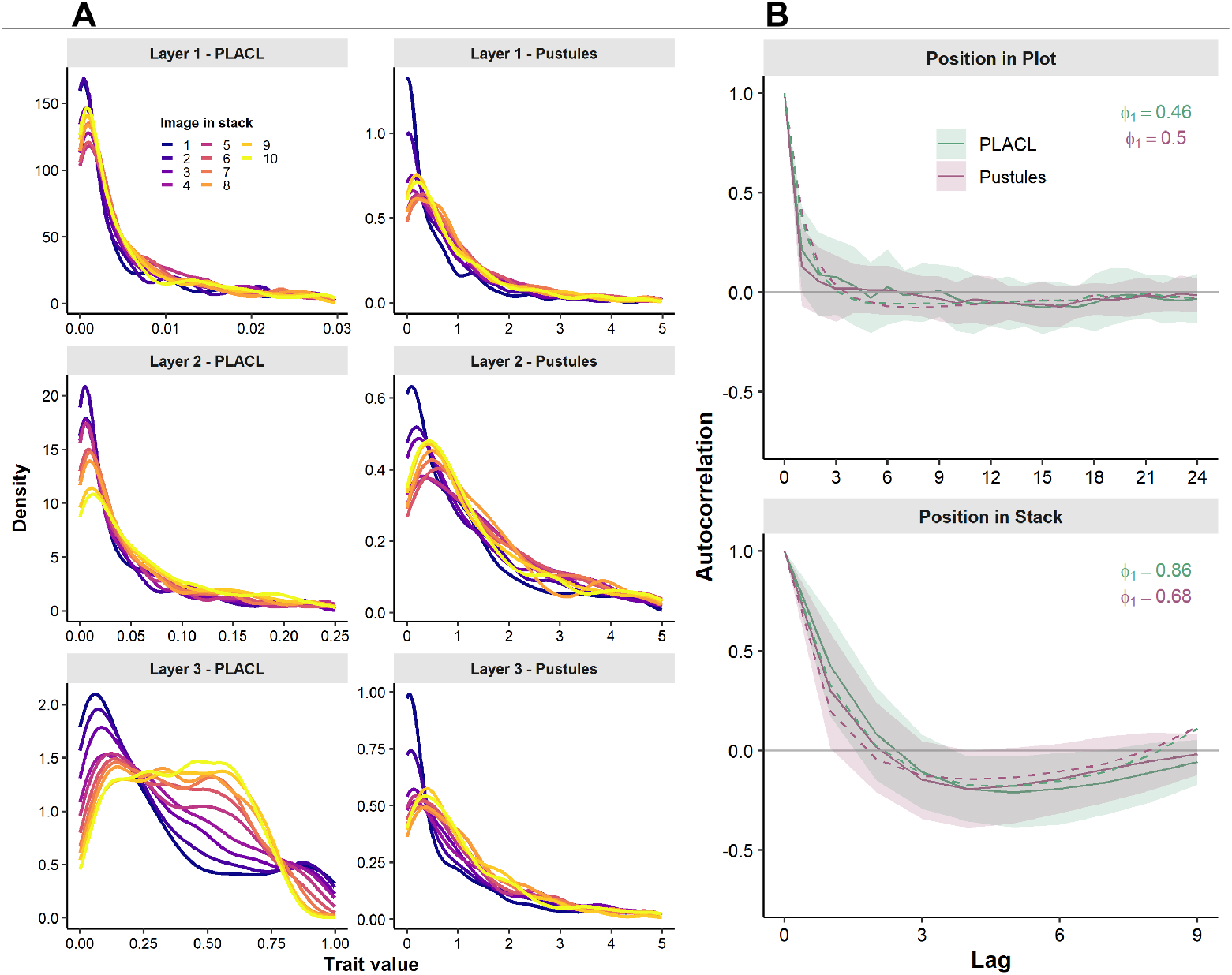
Autocorrelation patterns in trait values extracted from focal stacks and across plot positions. (A) Distribution of PLACL and rust pustule density across position-in-stack. Images 1 and 10 correspond to the first and last images in each stack, respectively. Values represent per–leaf layer means across all positions and plots. Note the variable x-axis scale varies for improved visualization; (B) Sample-based and model-based estimates of serial autocorrelation in focal stacks and across plot positions. Solid lines and shaded ribbons show the overall mean and standard deviation of the sample-based autocorrelation functions, while dashed lines indicate the model-based estimates. The parameter *ϕ*_1_ denotes the estimated AR(1) autoregressive coefficient, expressed on the transformed scale (logit for PLACL and log for rust pustule density).

On average, stack-level estimated autocorrelation at lag-1 was approximately *ρ*_1_ = 0.43 for PLACL and *ρ*_1_ = 0.30 for rust pustule density. Autocorrelation dropped to *ρ*_2_ = 0.09 and *ρ*_2_ = 0.02 at lag-2, subsequently becoming slightly negative at lags ≥ 3, to approach asymptotically *ρ* = 0 at lags ≥ 8 (Figure 6B, upper panel). Sample-based autocorrelation estimates were noisy, as expected due to small sample size (Supplementary Figure S1A), but trends were highly consistent across genotypes, genotype groups according to flag leaf angle, and vertical leaf layers (Supplementary Figure S1B-D). Sample-based autocorrelation estimates for position-in-plot were lower, estimated at *ρ*_1_ = 0.21 and *ρ*_1_ = 0.13 for PLACL and rust, respectively, then approaching asymptotically *ρ* = 0 at lags ≥ 2; Figure 6B, lower panel). This pattern was also highly consistent across plots and leaf layers (not shown).

The model autoregressive coefficients were high for position-in-stack (*ϕ*_1_ = 0.86 and *ϕ*_1_ = 0.68 for PLACL and rust density, respectively; Figure 6B, upper panel), but lower for position-in-plot and similar for both traits (*ϕ*_1_ = 0.46 and *ϕ*_1_ = 0.50 for PLACL and rust density, respectively; Figure 6B, lower panel). These autoregressive coefficients represent correlations on the transformed scale. Given our interest in the original scale, the estimated model parameters were used to simulate data, which were then back-transformed to the original scale to obtain a model-based estimate of autocorrelation on the original scale. This yielded lower estimates of autocorrelation compared with the overall means of the sample-based estimates for position-in-stack (*ρ*_1_ = 0.33 and *ρ*_1_ = 0.20 for PLACL and rust, respectively), whereas the opposite was the case for position-in-plot, where lag-1 autocorrelation was estimated at *ρ*_1_ = 0.38 and *ρ*_1_ = 0.40 for PLACL and rust, respectively; Figure 6B).

Despite the differences between sample-based and model-based autocorrelation decay patterns (Figure 6B), the estimated effective sample sizes were similar across both approaches. Somewhat surprisingly, ESF was estimated to be very similar for images in focal stacks and stacks across plot positions (Table 1). In other words, a focal stack consisting of 10 images represented approximately 2.5 independent measurements; 10 positions per plot with the realized pitch of approximately 14 cm also represented roughly 2.5 independent measurements.

**Table 1:** Effective sample fraction (ESF) by level, method, and trait.

| Level | Method | Trait | ESF |
| --- | --- | --- | --- |
| position-in-plot | sample-based | PLACL | 0.238 |
|  |  | Rust | 0.262 |
|  | model-based | PLACL | 0.214 |
|  |  | Rust | 0.211 |
| position-in-stack | sample-based | PLACL | 0.233 |
|  |  | Rust | 0.253 |
|  | model-based | PLACL | 0.253 |
|  |  | Rust | 0.273 |

## 5 Discussion

Although well known to photographers and widely used in microscopy, focus bracketing has, to our knowledge, not been employed for field phenotyping up until now. Typically, imaging systems for field phenotyping are designed to maximize depth-of-field by optical means, such as using a small aperture, and by capturing objects at distances where the natural depth-of-field is already sufficiently large. In addition, some focus blur can often be tolerated because the full image resolution may not be required or objects to be detected or segmented may be far larger than the blur kernel (e.g., [3]). Here, our objective was to combine symptom-level diagnosis based on miniature diagnostic features with canopy-level quantification of disease, which requires a very high spatial resolution and sufficient leaf area to be analyzed at the same time. The resulting trade-off between precision and variance of individual measurements represents a novel challenge in field phenotyping that, to the best of our knowledge, has not been addressed previously.

Focus bracketing as used here increased the effective sample size in terms of leaf area analyzed per measurement position by a factor of approximately 2.5, without compromising image quality, and at negligible marginal cost. While a single image yielded an analyzed area of 19.7cm^2^ on average, comparable to the average area of a sampled fully developed wheat leaf (17cm^2^ on average; [18]), focus bracketing increased this area to an equivalent of almost three sampled leaves. Somewhat surprisingly, our analysis demonstrated that consecutive images in focal stacks provided similar information as two separate images taken at close-by locations within a plot (Table 1; Figure 6). Where available, repeat measurements in previous studies were typically related to distinct biological entities, such as sampled leaves. In contrast, our approach evaluates contiguous patches of canopy captured within the camera’s field of view and located on the current focal plane, without reference to specific organs or plants. This enables the analysis of parts of leaves that may belong to different plants, arguably increasing the representativeness of the measurement for the experimental unit, as multiple independent leaves from separate plants are sampled. Thus, focus bracketing represents a promising approach to decrease the variance of disease severity estimates by increasing the effective leaf area that supports that estimate.

Previous studies typically relied on single measurements of disease per experimental unit [14, 26, 30, 15, 1], or on the mean of multiple measurements at each time point [34, 19, 1, 38] for downstream analyses. In some cases, within-plot variability of repeat observations was taken into account by including measures of variance as weights in downstream statistical analyses such as the estimation of heritability, spatially corrected plot values, or genotypic effects (e.g., [2]). In addition, several reports have already qualitatively described systematic trends in the precision and variance of disease severity assessments [8, 19]: for example, it has been established that the highest variance in individual measurements is observed around a mean severity of 0.5. This phenomenon has been independently observed in visual assessments [7], analysis of detached leaves [19], and in our own data (see Figures 3 and 4). However, an in-depth quantitative analysis of such relationships has not been performed.

Modeling disease severity estimates at the plot-level using the Beta distribution enabled us to derive estimates of the number of measurements required to achieve a given precision. The Beta distribution provided a robust framework to parameterize observed distributions across plots exhibiting a wide range of disease severity (Figure 4). In addition, it effectively captured the relationship between the mean and variance of disease severity. For example, to estimate the mean PLACL with a precision of *±*0.05 at a confidence level of 90%, between 1 and 84 samples are required, depending on the underlying distribution (see Figure 4). This has important practical implications, as it suggests that measurement intensity should be dynamically adapted to maintain a certain level of precision under evolving epidemics.

An assessment performed at an ideal time point where genotypes differ maximally with respect to disease intensity reduces precision requirements. However, identifying a single, ideal assessment time point may be impractical or not optimal in many situations: It would miss prominent genotype rank changes over the course of the season (e.g., [19]) and result in confounding with phenology. In contrast, conducting measurements at multiple time points to track epidemic development provides information about quantitative resistance that is, in many cases, essentially expressed in terms of its effects on canopy-level epidemics [36].

The leaf area analyzed per plot is directly related to the precision of disease severity estimates; consequently, maximizing the evaluated leaf area is a key objective. Because the proposed imaging approach does not require direct manual interaction with the canopy, it is, in principle, amenable to implementation on autonomous ground-based phenotyping platforms. Several challenges must be addressed to enable such a transfer, including (i) dynamic adjustment of camera position and orientation to accommodate differences in cultivar height, morphology, and canopy architecture, (ii) increased robustness of the image processing workflow to scene variability arising from seasonal changes in illumination as well as phenological and growth-related changes. The adopted side-view perspective is also unlikely to be optimal. First, the prominent relationship we found between position-in-stack and estimated disease severity (Figure 6A) is likely due to the greater contribution of lower vertical canopy fractions to the in-focus fraction of images with increasing position-in-stack, which suggests a potential bias related to canopy density and architecture, although microclimate-related effects (e.g., border rows versus interior plot positions) may also have played a role. Second, the scannable area is largely restricted to border regions, which can introduce systematic bias in plot-based experiments. Third, the current approach does not allow to exploit depth information to localize disease symptoms along the vertical canopy gradient, an opportunity that could be realized by adopting a nadir or oblique top-down imaging geometry. Ideally, the entirety of the canopy would be imaged; this represents a plausible scenario for single rows that are often used for disease screening during pre-breeding or early breeding generations. Under such conditions, the dominant source of uncertainty in estimating disease severity would shift from measurement-related limitations toward biologically and environmentally driven variability in disease expression.

## 6 Acknowledgments

We thank L. Roth (ETH Crop Science) for a thorough review of the statistical analyses, and J. Alassimone (ETH Plant Pathology) for instructions and practical assistance in preparing the artificial pathogen inoculations. We gratefully acknowledge support from the Group of Crop Science at ETH Zürich, especially A. Walter for sharing field and human resources, S. Corrado for crop husbandry, and B. Herzog for assistance with seed preparation and management. Funding: This work was funded by ETH Zürich.

## Author contributions

RZ: Conceptualization, investigation, data curation, methodology, formal analysis, visualization, software, and writing - review and editing. BAM: Conceptualization, supervision, funding acquisition, and writing - review and editing. JA: Conceptualization, investigation, methodology, formal analysis, visualization, software, writing - original draft, writing - review and editing, supervision, and project administration.

## Competing interests

The authors declare that they have no competing interests.

## 7 Data Availability

Images and extracted trait data will be made available by the authors upon reasonable request, without reservation.

## A Supplementary Information

### A.1 Supplementary Figures

**Figure S1.**
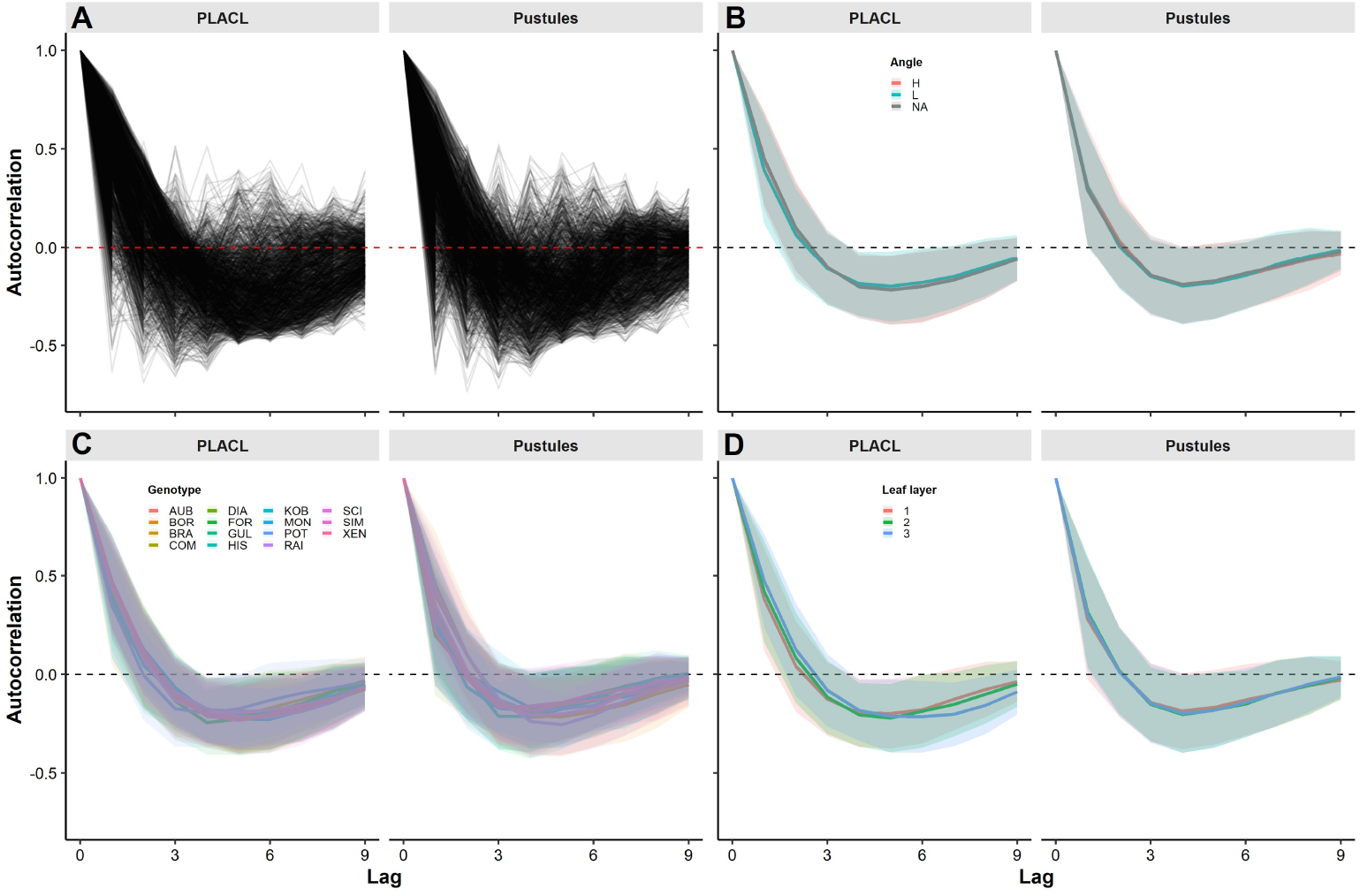
Sample-based autocorrelation of trait values across position-in-stack. (A) Individual sample-based autocorrelation function. (B-D) Mean and standard deviation by groups of cultivars: (B) according to flag leaf insertion angle (H: Large angle; L: small insertion angle); (C) according to cultivar; (D) according to leaf layer (1: Flag leaf; 2: Flag leaf minus one; 3: Flag leaf minus two

**Figure S2.**
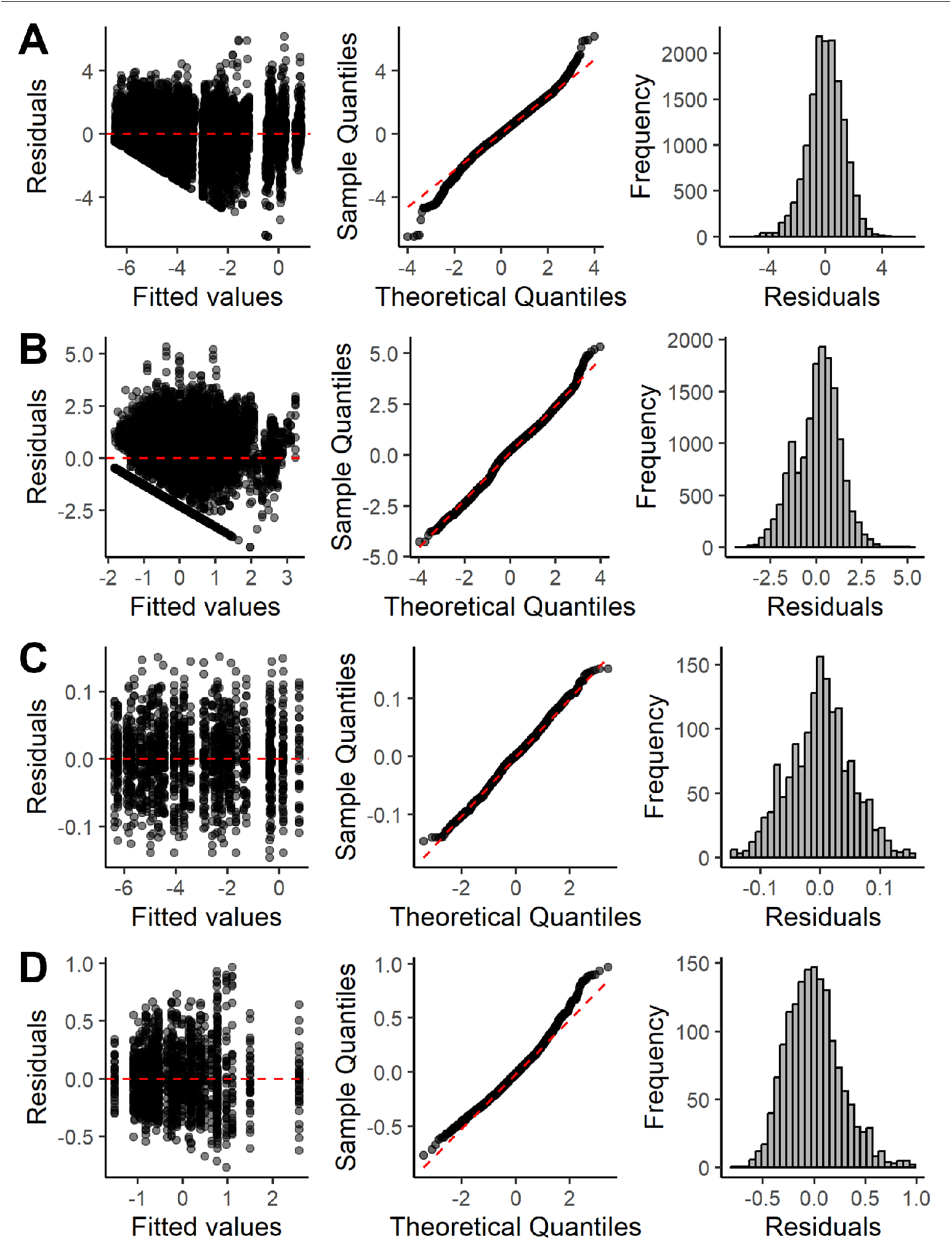
Standard diagnostics plots for the linear mixed effects models. (A) Residual plots for the position-in-stack autocorrelation model using logit-transformed PLACL values; (B) residual plots for the position-in-stack autocorrelation model using log-transformed rust pustule density values; (C) residual plots for the position-in-plot autocorrelation model using logit-transformed PLACL values; (D) residual plots for the position-in-plot autocorrelation model using log-transformed rust pustule density values.

## Notes

### Competing Interest Statement

The authors have declared no competing interest.

